# Lipid suppressed and tissue-fraction corrected metabolic distributions in human central brain structures using 2D ^1^H Magnetic Resonance Spectroscopic Imaging at 7 tesla

**DOI:** 10.1101/2020.06.09.142646

**Authors:** Alex A. Bhogal, Tommy A.A. Broeders, Lisan Morsinkhof, Mirte Edens, Sahar Nassirpour, Paul Chang, Dennis Klomp, Christiaan H. Vinkers, Jannie P. Wijnen

**Affiliations:** Department of Radiology, University Medical Center Utrecht, Utrecht, The Netherlands; Technical Medicine, University of Twente, Enchede, The Netherlands; MR Shim GmbH, Reutlingen, Germany; Department of Psychiatry, Brain Center Rudolf Magnus, University Medical Center Utrecht, Utrecht, The Netherlands; Department of Anatomy & Neurosciences, Amsterdam UMC (location VU University Medical Center), Amsterdam, The Netherlands; Department of Psychiatry, Amsterdam UMC (location VU University Medical Center)/GGZ inGeest, Amsterdam, The Netherlands

**Keywords:** MRSI, metabolic imaging, 7 tesla, proton spectroscopy, glutamate

## Abstract

Magnetic resonance spectroscopic imaging (MRSI) has the potential to add a layer of understanding of the neurobiological mechanisms underlying brain diseases, disease progression, and treatment efficacy. Limitations related to metabolite fitting of low SNR data, signal variations due to partial volume effects, acquisition and extra-cranial lipid artefacts, along with clinically relevant aspects such as scan-time constraints, are among the factors that hinder the widespread implementation of *in vivo* MRSI. The aim of this work was to address these factors and to develop an acquisition, reconstruction and post-processing pipeline to derive lipid suppressed metabolite values based on Free Induction Decay (FID-MRSI) measurements made using a 7 tesla MR scanner. Anatomical images were used to perform high-resolution (1mm^3^) partial-volume correction to account for grey matter, white matter and cerebral-spinal fluid signal contributions. Implementation of automatic quality control thresholds and normalization of metabolic maps from 23 subjects to the MNI standard atlas facilitated the creation of high-resolution average metabolite maps of several clinically relevant metabolites in central brain regions, while accounting for macromolecular distributions. Reported metabolite values include glutamate, choline, (phospo)creatine, myo-inositol, glutathione, N-acetyl aspartyl glutamate(and glutamine) and N-acetyl aspartate. MNI-registered average metabolite maps facilitate group-based analysis; thus offering the possibility to mitigate uncertainty in variable MRSI.

## 1.0 INTRODUCTION

Magnetic resonance spectroscopy (MRS) is a technique that is used to acquire in-vivo brain tissue metabolite signals from single, rather large, preselected cubic volumes that typically occupy approximately 8 cm^3^. The highly localized nature of this acquisition sharply contrasts with the ubiquitous presence of metabolites across the entire brain. Advances in MRI hardware have encouraged the development of advanced spectroscopic techniques (Magnetic Resonance Spectroscopic Imaging (MRSI) to measure neurochemical distributions at increasingly high spatial resolutions. In contrast to single-voxel MRS, MRSI is sensitive to diffuse and potentially heterogeneous changes in brain metabolites. As a result, MRSI has the potential to fundamentally enhance our understanding of *in vivo* neurochemical processes and provides a potential tool to explore cases in which pathophysiological origins of disease remain unclear or where standard structural MRI prove to be of limited diagnostic use. These new opportunities come partially as the result of the availability of ultra-high field (>=7 Tesla) MRI scanners that provide higher peak signal-to-noise ratios (SNR), as well as increased spectral dispersion to facilitate separation of overlapping metabolite resonances (for an overview see [1]). Higher SNR improves the detection capacity resonances such as glutamate, and/or can be translated into increased MRSI resolution that can range between 2-5 mm^2^ in-plane with slice thickness in the order of 10-12 mm at 9.4[2] and 7 Tesla (7T) [3], respectively.

Contemporary studies reporting metabolite distributions measured using ultra-high field (7T+) MR scanners have mainly demonstrated ‘proof of concept’ or methodological developments in specific areas such as acquisition[4-7] or data reconstruction [8-10]. The translation of these techniques to the clinical environment remains hindered by sensitivity to artefacts emanating from unsuppressed water resonances (at 4.7ppm), signal contamination by extra-cranial lipids (0.9-1.3ppm) [11], and artefacts caused by inhomogeneous B_1_ and B_0_ fields. Ongoing research focuses on overcoming some of these challenges.

Various approaches to address the issue of inhomogeneous B_0_ at ultra-high field strength exist, including slice-based shimming when requisite hardware is available [12], or post-processing methods involving field-map based B_0_ correction [13]. Overwhelming water signals can be suppressed using saturation pre-pulses [14-16] or inversion-recovery based nulling of the water signal before excitation [17]. Lipid signal suppression is possible via outer volume saturation (OVS) slabs [18], however OVS solutions necessitate long repetition times (TR) due to restrictive specific absorption rate (SAR) limits set on radio frequency (RF) pulses. The resulting prohibitively long scan-times can be mitigated with the addition of accelerated data acquisition strategies [4, 6, 19], replacement of OVS with post-processing based lipid signal removal techniques [20] or advanced spectral-spatial RF pulses [21]. These methods are not without their respective trade-offs and potentially introduce unwanted artefacts or practical limitations that may restrict implementation in larger-scale studies. Alternative hardware-based approaches can also forgo the need for the SAR-intensive OVS pulses through direct suppression of undesired lipid signals [22, 23]. However, such technologies can lead to collateral suppression of tissue metabolite signals that are of interest.

The aim of this study was to implement a novel combination of MRSI acquisition, reconstruction and post-processing methods available at our institution in order to derive lipid-suppressed spectra at 7T and associated metabolite values for central brain structures. We used a 7 tesla MR scanner to acquire high resolution, short echo-time (TE) MRSI data along with an external crusher coil [22] for hardware based extra-cranial lipid signal suppression. Water suppression was achieved using a shortened chemical shift selective (CHESS) sequence with subject specific spectral-spatial (tailored) B_1_-insensitive suppression pulses [14]. Advanced reconstruction and post-processing techniques were used in conjunction with stringent data filtering based on several quality assurance (QA) metrics[24] including linewidth, SNR, Cramér-Rao Lower Bounds (CRLB) and the Lipid-to-total-Creatine ratio. High-resolution anatomical images facilitated a point-spread function adjusted[11, 25] partial volume correction (PVC) method that accounted for grey matter (GM) and white matter (WM) signal contributions, along with the more commonly applied correction for cerebral-spinal fluid (CSF). Finally, registration of resulting metabolic maps to the MNI152 standard atlas facilitated the creation of high-resolution average metabolite maps, in which macromolecular signal contributions were also accounted for [26].

## 2.0 METHODS

This study was approved by the medical research ethics committee of University Medical Center Utrecht and written informed consent was obtained from all subjects. The experiments were performed according to the guidelines and regulations of the WMO (Wet Medisch Wetenschappelijk Onderzoek). Data were acquired in 23 healthy volunteers that were recruited from the general population (age 23 ± 5 years, 9 females) using a 7 tesla MRI scanner (Phillips, Best, NL) equipped with a dual-transmit head coil in combination with a 32 channel receive coil (Nova Medical, Wilmington Ma, USA). Second order image-based shimming was performed for all acquisitions.

### 2.1 MRSI data acquisition

Suppression of extra-cranial lipid signals, and therefore, significant lipid contamination artefacts [11], was achieved using an external crusher coil [22] driven by an external amplifier capable of delivering +/- 10A of current. To ensure patient safety and prevent overheating of the coil windings, 1.5A fuse was installed between the amplifier and crusher coil. The amplifier ramp time is slow relative to the crusher coil pulse duration and the fuse burns when the integral of the current over this period exceeds 1.5A. Image-based calibration of the crusher coil amplified settings are described in supplementary figure 2. MRSI data were acquired using a slice-selective, free-induction decay sequence [3] with the following parameters: acquisition delay/TR = 2.5/300 ms, FOV = 220×220 mm, acquisition matrix = 44×44, resolution 5×5×10 mm^3^, BW = 3000 Hz, samples = 512, signal averages = 2, elliptical k-space shuttering, scan duration = 10 m 59 s, tailored spiral in-out spectral-spatial water suppression pulses [14], flip angle = 35°. Water unsuppressed MRSI data were acquired for zeroth order phase and eddy current correction with adapted parameters: acquisition matrix = 22×22, resolution 10×10×10mm^3^, signal averages = 1, scan duration = 1m 54s. Two adjacent 10mm MRSI slices (20mm slab) were acquired axially to intersect deep GM nuclei located adjacent to the ventricles while maximizing the amount of tissue acquired through the occipital lobe. This region is of interest since many MRSI studies performed at UHF thus far have focused primarily on brain regions above the ventricles, whereas neuropsychiatric disorders (schizophrenia or psychosis, but also obsessive compulsive disorder) are believed to have origins, at least partly, in deep gray matter structures[27].

In a single volunteer, a double inversion recovery sequence in combination with lipid signal suppression using the crusher coil was used to acquire a measured macromolecular baseline signal [26]. The slice was planned above the ventricles to minimize artefacts relating to unsuppressed water signal and B_0_ inhomogeneity; scan parameters were: TE/TR = 2.5/1000 ms, FOV = 220×220 mm^2^ resolution 30×30×10 mm^3^, BW = 6000 Hz, samples = 512, elliptical k-space, signal averages = 30, scan duration = 11m 28s. The inversion pulses were inserted within a VAPOR water suppression sequence with the first pulse at (TI_1_) 870ms before excitation, and the second, (TI_2_) 296 ms before excitation. The bandwidth of the inversion pulse was 6.7 ppm (2 kHz) with an offset of -3.7 ppm (−1110 Hz) and center pulse at 1 ppm. Thus, metabolite nulling was covered up to 4.3 ppm and did not affect the water resonance at 4.7 ppm. Spectral quality was evaluated visually and a total of 6 voxels were isolated and averaged to generate the metabolite and lipid suppressed MM signal. A series of 9 Gaussian functions were fit to the MM signal using AMARES (jMRUI version 3.0). To minimize the degrees of freedom associated with overlapping resonances, certain MM lines were grouped (see supplementary figure 3). Chemical shift values were as follows: MM_09 (0.94 ppm), MM_12_14 (1.29 ppm & 1.42 ppm), MM_17_20 (1.79 ppm & 2.04 ppm), MM_23 (2.31 ppm), MM_27 (2.75 ppm), MM_30_32 (3.02 ppm & 3.25 ppm).

### 2.2 Imaging

A 3D Turbo Field Echo (TFE) scan (TE/TR = 2.89/8ms, resolution = 1mm isotropic, FOV = 220×220×200 mm^3^, scan duration = 6min 51sec, flip angle = 6 degrees) along with a shimmed dual echo GRE B_0_ map (delta TE/TR = 2.35/5.12 ms, FOV = 220×220×30 mm^3^, acquisition matrix = 176×176, resolution = 1.25 × 1.25 × 10mm^3^, slices = 3, scan duration, 1m 6s) and 2D multi-slice T1w Fast Field Echo (FFE) image (TE/TR = 4.22/200 ms, FOV = 220×220×30 mm^3^, acquisition matrix = 176×176, resolution = 1.25 × 1.25 × 10mm^3^, slices = 3, scan duration, 72 s) were acquired. A reconstruction of 20 slices, each with 1mm thickness (termed ‘***slice***’), occupying the same volume as the 20 mm MRSI slab was made on the scanner console based on the 3D-TFE scan.

### 2.3 Reconstruction and Post-Processing

Reconstruction and processing was performed using an automatic Python-based MRSI analysis pipeline that applied the following steps: up-sampling of the MRSI acquisition grid by an in-plane factor of 4×4 such that a single MRSI voxel became 16 voxels to match with the shimmed B_0_ map. Reconstruction and channel combination using coil sensitivity profiles derived from the 2D T1w FFE image (ESPIRiT method described in [28]) followed by over-discretized B_0_ correction based on the shimmed B_0_ map using algorithms described in: [11, 13]. After reconstruction, MRSI data were down-sampled to the acquired 44×44 grid using an optimized Gaussian spatial response function. These steps served to reduce near- and far-reaching voxel bleeding from residual extra-cranial lipid signals (for details see: [11, 25]), improve overall SNR via over-discretized noise-decorrelation and enhance spectral linewidth through over-discretized spectral re-alignment [13]. Eddy current correction [29] and zero-order phase correction were performed using the unsuppressed water data and residual water signal was removed using the Hankel Lanczos (HLSVD) method. Automatic first-order phase correction, was achieved using a backward Yule-Walker linear prediction autoregressive algorithm on each free-induction decay (FID) signal. Total reconstruction and processing time was approximately 5 minutes for each dataset.

### 2.4 LCModel Fit & Metabolite Map Generation

1H MR spectral profiles were simulated for each relevant compound by NMRSIM (v. 4.6.a. Bruker Biospin, Billerica, MA) based on a pulse acquire sequence with an TE of 0.001ms (first order phase correction performed during reconstruction outlined in 2.3). The basis set included tCho: glycerophosphocholine (GPC) and choline (Cho), phosphocholine (PC), tCr: creatine (Cr) and phosphocreatine (PCr), Glx: glutamate (Glu) an glutamine (Gln), taurine (Tau), myo-inositol (mI), glycine (Gly), glucose (Glc), tNAA: N-acetylaspartate(glutamate) (NAA(G)), gamma-aminobutyrate (GABA), aspartate (Asp), glutathione (GSH), lactate (Lac), succinate (Suc), guani-doacetate (Gua), scyllo-Inositol (Scyllo), and acetate (Ace). Chemical shift coupling constants reported in Govindaraju et. al [30] were implemented. All simulated spectral profiles together with the measured macromolecular signals served as the basis set to fit measured in vivo spectra using LCModel [31] (version 6.3-1K). The relative linewidths were constrained to the linewidth of MM09, while amplitudes of MM components were constrained in cases where they were grouped (i.e. these peaks were bound to one another: see supplementary figure 3). Lipid resonances (0.9 ppm, 1.3 ppm (Lip13ab), 2.0 ppm) were simulated by LCModel. All spectra were fit between 0.2 and 4.0 ppm. The complete basis set and LCModel control parameters are included in the supplementary material.

Fitted metabolite values along with quality assurance (QA) data consisting of Cramer Rao Lower bounds (CRLB - %Standard Deviation), SNR and full-width at half maximum (FWHM) outputs were converted into 2D maps using custom Matlab scripts. QA maps formed the basis for binary filters that were applied to discard metabolite voxels which did not meet the following criteria: SNR > 3, FWHM < 0.15 ppm, Lip13ab/tCr < 2 or CRLB < 50%. CRLB filters based on the LCModel output were applied on a per-metabolite/MM basis. Since maps were expressed as a ratio with tCr, voxels having a CRLB higher than 50% in the tCr map were removed from all metabolite maps. A caveat regarding the use of CRLB for QA is that it may be ill-suited in cases where metabolite values are expected to change due to disease (for analysis see [32]).

Metabolic and QA maps generated from the LCModel output were regridded to the same in-plane/in-slice resolution as the *slice* anatomical image to mirror the reconstruction made from the 3D T1w anatomical scan (i.e. resolution from 5×5×10mm^3^ to 1mm^3^; matrix from 44×44×1 to 220×220×10). This step simply subdivided a single MRSI voxel into 250 voxels using nearest-neighbor interpolation to enable high-resolution partial volume correction and the subsequent application of transformation matrices derived during registration of anatomical images to MNI space.

### 2.5 Partial Volume and T1 Correction

We extend the partial volume correction (PVC) typically done to account for CSF fraction in MRS voxels to include corrections for GM and WM tissue fractions (schematically outlined in supplementary figure 1). To achieve this, MRSI voxel intensities were expressed as a weighted sum of pure tissue contribution. Weighting coefficients were derived based on GM, WM and CSF segmentations (FSL: FAST) that provided the tissue’s fractional volume within the voxel. For a corollary, the reader is encouraged to see similar PVC methods that have been applied to perfusion maps derived from Arterial Spin Labelling experiments [33] and examples of spectroscopic reports reporting brain metabolite values relative to tissue fraction [34, 35]. Based on prior knowledge regarding the MRSI point spread function (PSF) and Gaussian response function applied during reconstruction (see section 2.3 and [11, 13]), anatomical segmentations were convolved with an appropriate PSF to mimic the signal bleeding effects inherent to the MRSI acquisition (see figure 1B and methods in [25]). For generation of the PSF, the k-space shutter applied during acquisition, along with the acquisition matrix of the anatomical data used for PVC were taken into account. The steps used to generate the PSF are elaborated on in supplementary figure 4. The zero-padding associated with the elliptical k-space shutter and over-discrete reconstruction yielded a PSF with a 8.75 mm FWHM. Using this information, metabolite values within a voxel were then redistributed (within the same occupied volume) based on point-spread adjusted tissue fraction and pure tissue metabolite values (see supplementary figure 1). Due to low metabolite abundance, the CSF contribution to the total metabolite signal was considered to be negligible. Pure tissue contributions were estimated by linear regression of measured metabolite values against the normalized tissue fraction of the voxel (GM fraction divided by the sum of the GM and WM fractions) and extrapolating to a GM fraction of one (pure GM) or zero (pure WM)[35, 36]: see supplementary figure 1. Partial volume corrected maps were corrected for steady-state T1 relaxation effects based on tissue specific T1 values reported at 7T by Xin et. al [37]. T1 values are provided in supplementary table 1.

**Figure 1A:**
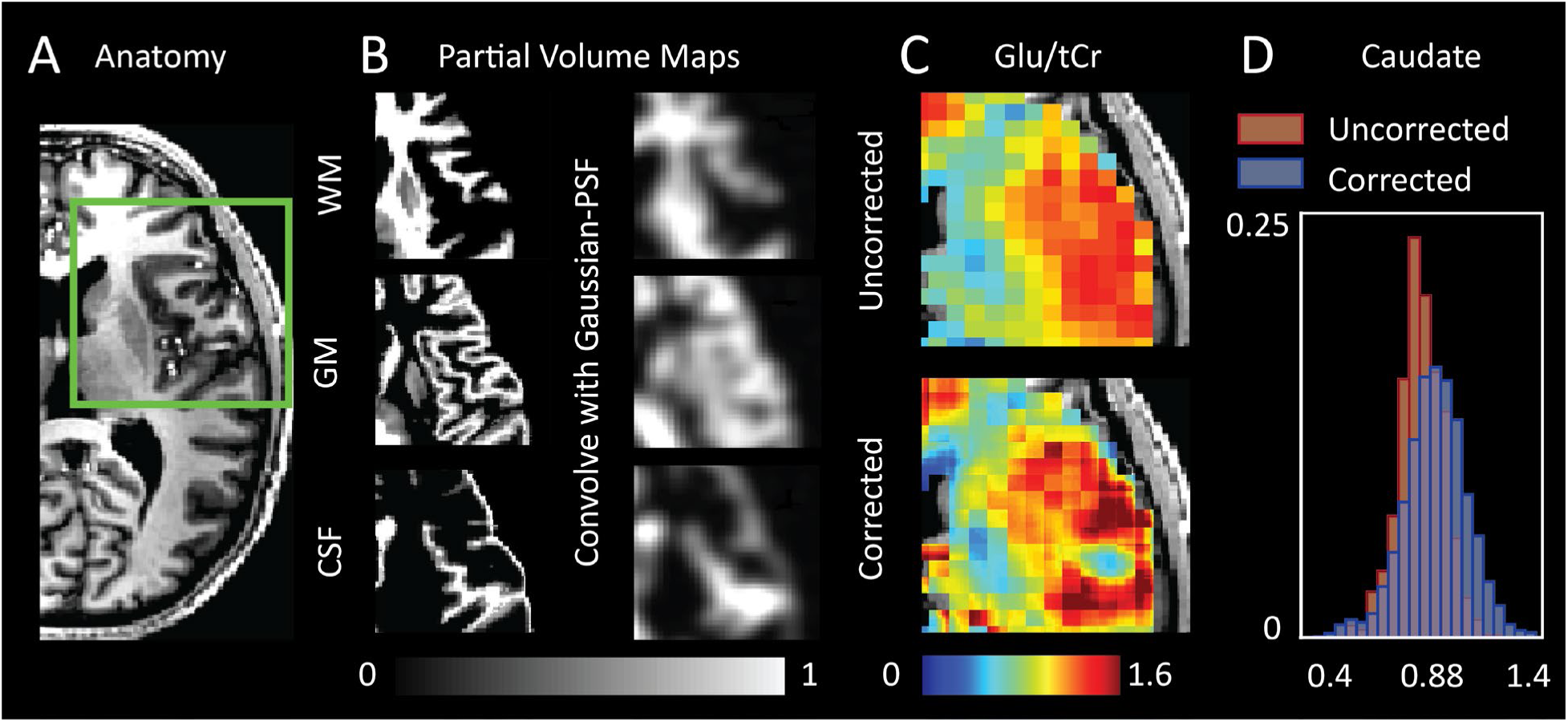
T1-weighted anatomical scan; 1B: GM, WM and CSF segmentations are convolved with an appropriate point-spread function to account for the signal dispersion (‘voxel bleeding’) effects associated with reconstructing lower resolution MRSI data; 1C: PSF-adjusted segmentations are used for partial volume correction of metabolic data. A map of Glu/tCr for a single subject before and after partial volume and T1 correction is shown; 1D: Example probability histogram derived from MNI-averaged Glu/tCr map in the Caudate Nucleus before and after PVC. PVC results in an increased standard deviation due to signal dispersion throughout voxels based on partial volume contributions, as well as a shift to higher mean Glu/tCr in deeper GM nuclei consistent with the removal of partial volume effects with WM at tissue border-zones.

### 2.6 Normalization of MRSI data to standard space (see figure 2)

Partial-volume corrected metabolite maps were normalized to MNI space by applying three transformation matrices generated using the FSL software library [38]. The first transformation (T_X1_) was derived from the rigid registration (FSL: FLIRT [39]) of the reconstructed *slice* image to the 3D T1w scan. Next, brain extracted anatomical images (optiBET shell script [40]) were registered to the 1mm MNI152 atlas via rigid (FLIRT) and then non-linear transformations (FSL: FNIRT[41]) to generate T_X2_ and T_X3_, respectively. Finally, transformations were applied, in order, to create MNI-registered metabolite and QA maps. These maps were averaged and the data density was calculated on a voxel-wise basis. To examine regional metabolite values, MNI masks were chosen based on their spatial correspondence with the metabolic data contained in the average maps; GM segmentations: accumbens, caudate, pallidium, putamen, thalamus, intracalcarine (IC) cortex, insular cortex and total cerebral cortex; white matter (WM) segmentations: corpus callosum, thalamic radiation and total white matter.

**Figure 2.**
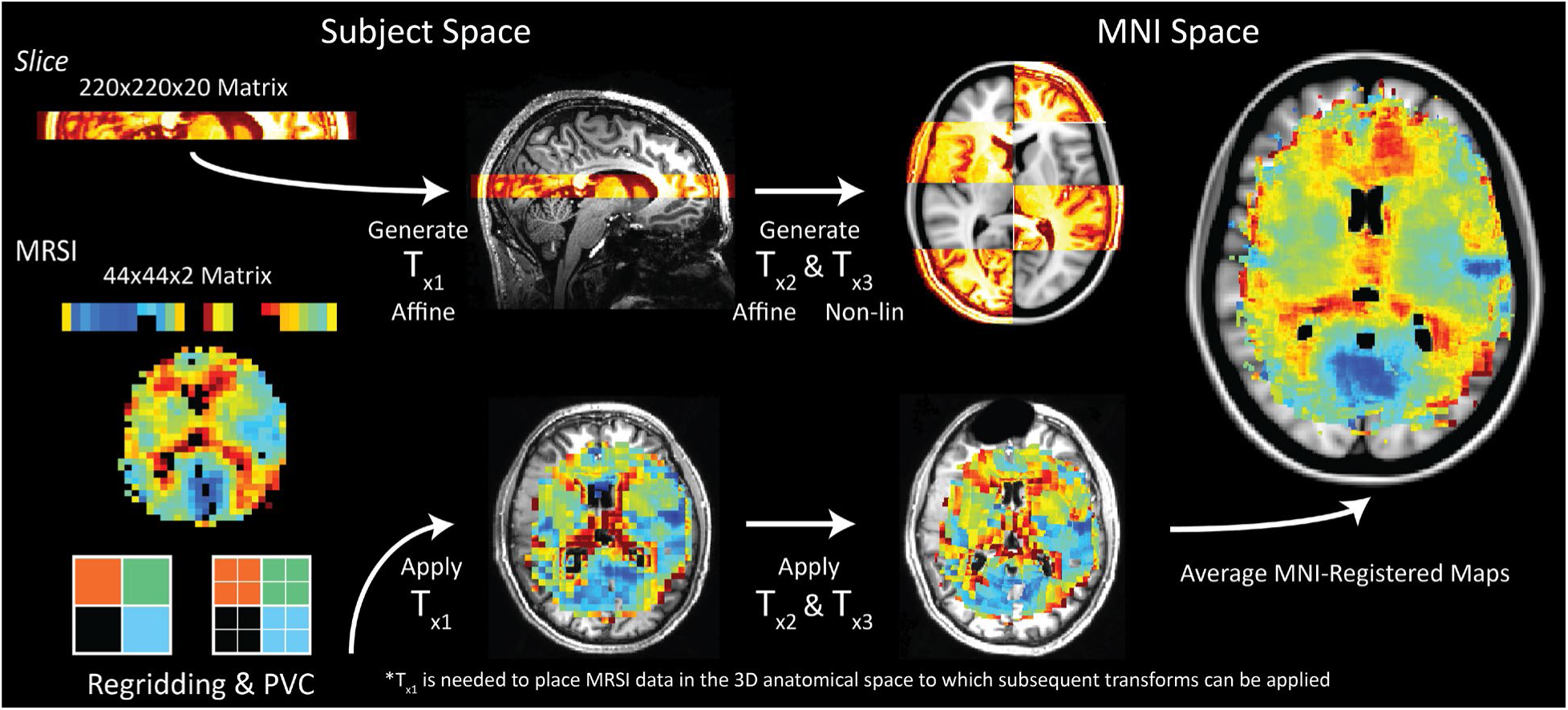
TOP: An anatomical ‘slice’ is reconstructed on the scanner console from the whole-brain 3D T1-weighted anatomical scan. This slice occupies the same volume as the MRSI acquisition. The slice is registered back to subject space (T1w image) using an affine transformation. The T1w image is then registered to MNI space via affine and non-linear transformations; 2-BOTTOM: Metabolic maps (unsmoothed tCho/tCr from a single subject is shown) are resampled to the same resolution as the reconstructed slice (nearest-neighbor interpolation). The transformation matrices (T_x1-3_) derived by registration of anatomical data from subject to MNI space are applied to the resampled metabolite maps. MNI-registered MRSI data is averaged in MNI space. The MNI-registered tCho map based on 23 subjects is shown (right).

## 3.0 RESULTS

Despite the ‘blurring’ of structural data resulting from convolution with the Gaussian-PSF (figure 1B), considerable differences between corrected and uncorrected metabolite maps were evident (figure 1C). Specifically, the effects of PVC were most obvious in regions containing CSF where metabolite signals were redistributed to neighboring tissues, as well as in regions containing transitions between GM and WM. The later effect was most notable for metabolites such as Glu/Glx and tCho, that are known to show regional variability[5, 42].

An indication of the amount of data removed for various QA filters is provided in supplementary table 2. Due to asymmetries in the lipid suppression gradient, cortical metabolite signals subject to the suppression field were consistently filtered out during the QA process. The contrast in quality between spectra originating in ‘over crushed’ versus ‘optimally crushed’ cortical regions are shown in figure 3. Over-crushed cortical regions were generally filtered out due to low SNR or high FWHM (figure 4).

**Figure 3.**
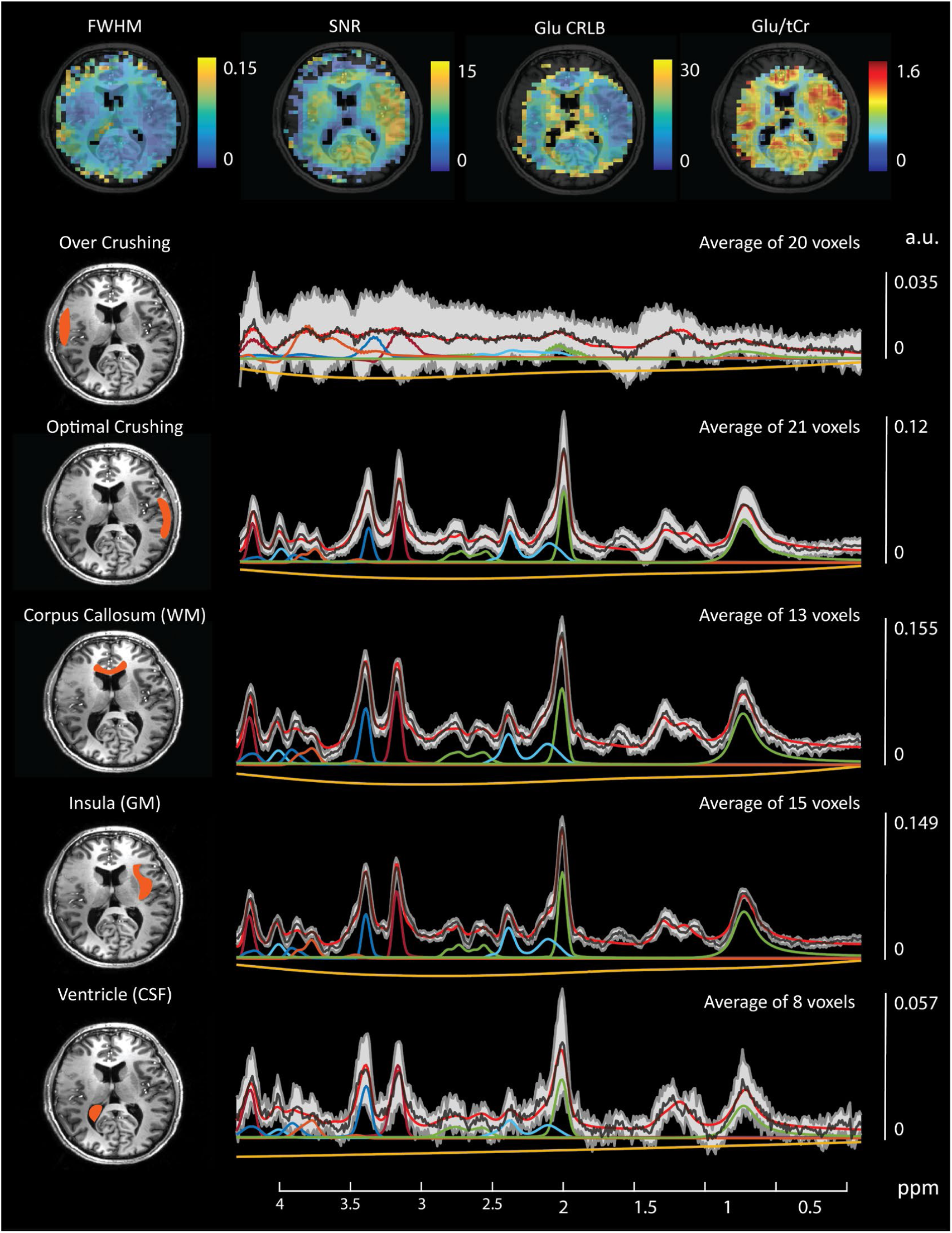
TOP: Example single-subject QA maps. Voxels with SNR values lower than 3, FWHM values higher than 0.15 ppm and CRLB values higher than 50% are removed. Thresholded QA maps are binarized and used to filter corresponding metabolite maps. A QA filtered Glu map is shown TOP-RIGHT. Overlay of metabolite maps with anatomical data allows precise delineation of ROIs for spectral plotting. Average spectra for example ROIs (left) are denoted by the black line with standard deviations defined by the grey shaded area. Corrections for inhomogeneous B_1_ were not performed so spectral signal intensity variations may show artificially large standard deviations (gray shaded area). Average fit values for selected metabolites are shown for each ROI: tNAA (green), Glx (light blue), tCr (red), tCho (dark blue), mI+Gly (orange), fit baseline (yellow), total fit (red), average signal (black).

**Figure 4:**
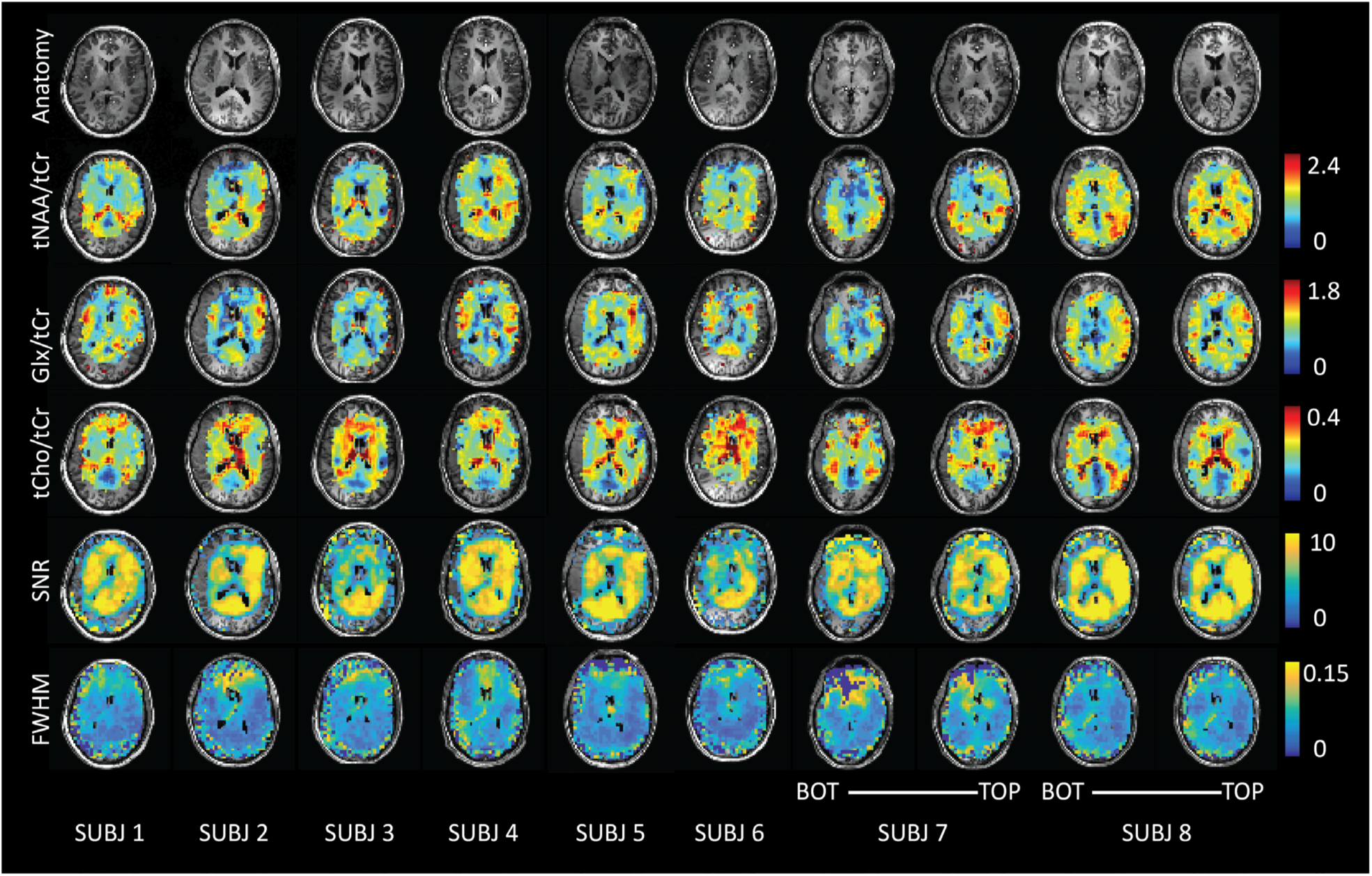
Partial volume corrected metabolite data of 8 subjects is shown for tNAA, Glu and tCho along with corresponding SNR and FWHM maps derived from the LCModel ‘miscellaneous output’. Both the bottom and top slices are shown for subject numbers 7 and 8.

Metabolite ratios are reported for applicable MNI segmentations in table 1. The known differences in Glu/tCr (or Glx/tCr) between GM and WM were reproduced[42]. Elevated Glu/tCr levels were measured in cortical structures (insula, IC and cerebral cortex), with reduced levels found in white matter and sub-cortical structures in line with results reported using single-voxel MRS at 7T [43]. These distributions are consistent with the role of Glu in energy production and neurotransmission. The opposite was observed in tCho/tCr maps, where levels in the occipital lobe, the insular and IC cortex and the cerebral cortex were likely influenced by partial volume effect with CSF. The levels of tCho/tCr reflect soluble membrane components [44] and since much of GM is occupied by microvasculature (and therefore blood), WM tCho/tCr levels were in line with expectations. In addition, the regional heterogeneity with elevated levels in frontal regions shown in our maps support single-voxel results presented by Pouwels et. al [45]. Levels of tNAA/tCr in the nucleus accumbens, caudate, pallidum and putamen were low as compared with the thalamus and cortical GM structures. High levels are expected in cortical structures since NAA, the dominant resonance around 2.0ppm, is considered a marker of viable neurons and neuronal density is highest in cortical GM regions. The difference in ratios between cortical/thalamic regions and basal ganglia may be explained by their varying physiological functions. The combined map of mI+Gly/tCr showed higher ratios in the corpus callosum and neighboring WM structures. Lower values were seen in cortical/insular structures. While ratios were higher in low-frontal regions of the brain, it is possible that this region was affected by inhomogeneous B_0_ around the nasal cavities, residual water signal around 4.7 ppm or low data density, and therefore greater uncertainty stemming from inter-subject variability. Nevertheless, GM versus WM differences in mI+Gly/tCr were apparent, which is consistent with previous single voxel experiments [46] and metabolic maps acquired at 9.4 T [2].

**Table 1.**
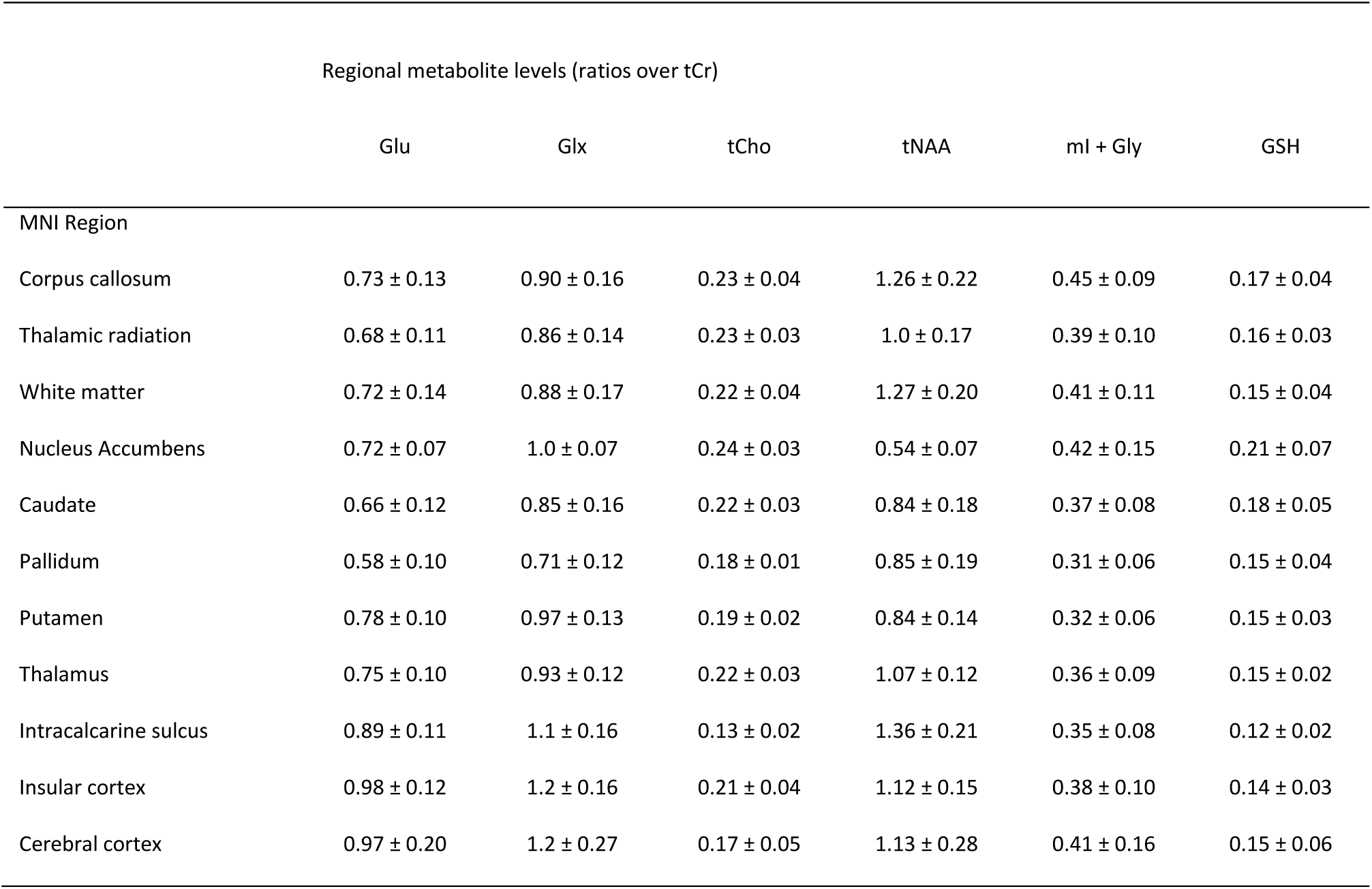

|  | Regional metabolite levels (ratios over tCr) |  |  |  |  |  |
| --- | --- | --- | --- | --- | --- | --- |
|  | Glu | Glx | tCho | tNAA | ml + Gly | GSH |
| MNI Region |  |  |  |  |  |  |
| Corpus callosum | 0.73 ± 0.13 | 0.90 ± 0.16 | 0.23 ± 0.04 | 1.26 ± 0.22 | 0.45 ± 0.09 | 0.17 ± 0.04 |
| Thalamic radiation | 0.68 ± 0.11 | 0.86 ± 0.14 | 0.23 ± 0.03 | 1.0 ± 0.17 | 0.39 ± 0.10 | 0.16 ± 0.03 |
| White matter | 0.72 ± 0.14 | 0.88 ± 0.17 | 0.22 ± 0.04 | 1.27 ± 0.20 | 0.41 ± 0.11 | 0.15 ± 0.04 |
| Nucleus Accumbens | 0.72 ± 0.07 | 1.0 ± 0.07 | 0.24 ± 0.03 | 0.54 ± 0.07 | 0.42 ± 0.15 | 0.21 ± 0.07 |
| Caudate | 0.66 ± 0.12 | 0.85 ± 0.16 | 0.22 ± 0.03 | 0.84 ± 0.18 | 0.37 ± 0.08 | 0.18 ± 0.05 |
| Pallidum | 0.58 ± 0.10 | 0.71 ± 0.12 | 0.18 ± 0.01 | 0.85 ± 0.19 | 0.31 ± 0.06 | 0.15 ± 0.04 |
| Putamen | 0.78 ± 0.10 | 0.97 ± 0.13 | 0.19 ± 0.02 | 0.84 ± 0.14 | 0.32 ± 0.06 | 0.15 ± 0.03 |
| Thalamus | 0.75 ± 0.10 | 0.93 ± 0.12 | 0.22 ± 0.03 | 1.07 ± 0.12 | 0.36 ± 0.09 | 0.15 ± 0.02 |
| Intracalcarine sulcus | 0.89 ± 0.11 | 1.1 ± 0.16 | 0.13 ± 0.02 | 1.36 ± 0.21 | 0.35 ± 0.08 | 0.12 ± 0.02 |
| Insular cortex | 0.98 ± 0.12 | 1.2 ± 0.16 | 0.21 ± 0.04 | 1.12 ± 0.15 | 0.38 ± 0.10 | 0.14 ± 0.03 |
| Cerebral cortex | 0.97 ± 0.20 | 1.2 ± 0.27 | 0.17 ± 0.05 | 1.13 ± 0.28 | 0.41 ± 0.16 | 0.15 ± 0.06 |

In addition to metabolic information, QA and data density information was evaluated in MNI space. As expected, an inverse spatial relationship was observed between SNR and linewidth FWHM (with regions of low SNR displaying broader linewidths; see figure 6). This was likely a result of inconsistencies in shimming, variations in the B_1_^+^ field or possible effects relating to the lipid suppression field. Reduced SNR with increased FWHM was observed at frontal regions of the brain, reflecting inhomogeneous B_0_ arising from susceptibility effects around the nasal cavities. This trend was consistent with higher CRLB values for tNAA at frontal regions of the brain. In comparison, CRLB values for Glx were higher along WM tracts (figure 6). Glx consists of low SNR metabolites with low WM abundance relative to GM. These factors contribute to greater fit uncertainty and relatively higher CRLB within WM regions. Density maps highlight increased information sampling at central regions of the brain consistent with acquisition of data at the level of the sub-cortical nuclei.

**Figure 5:**
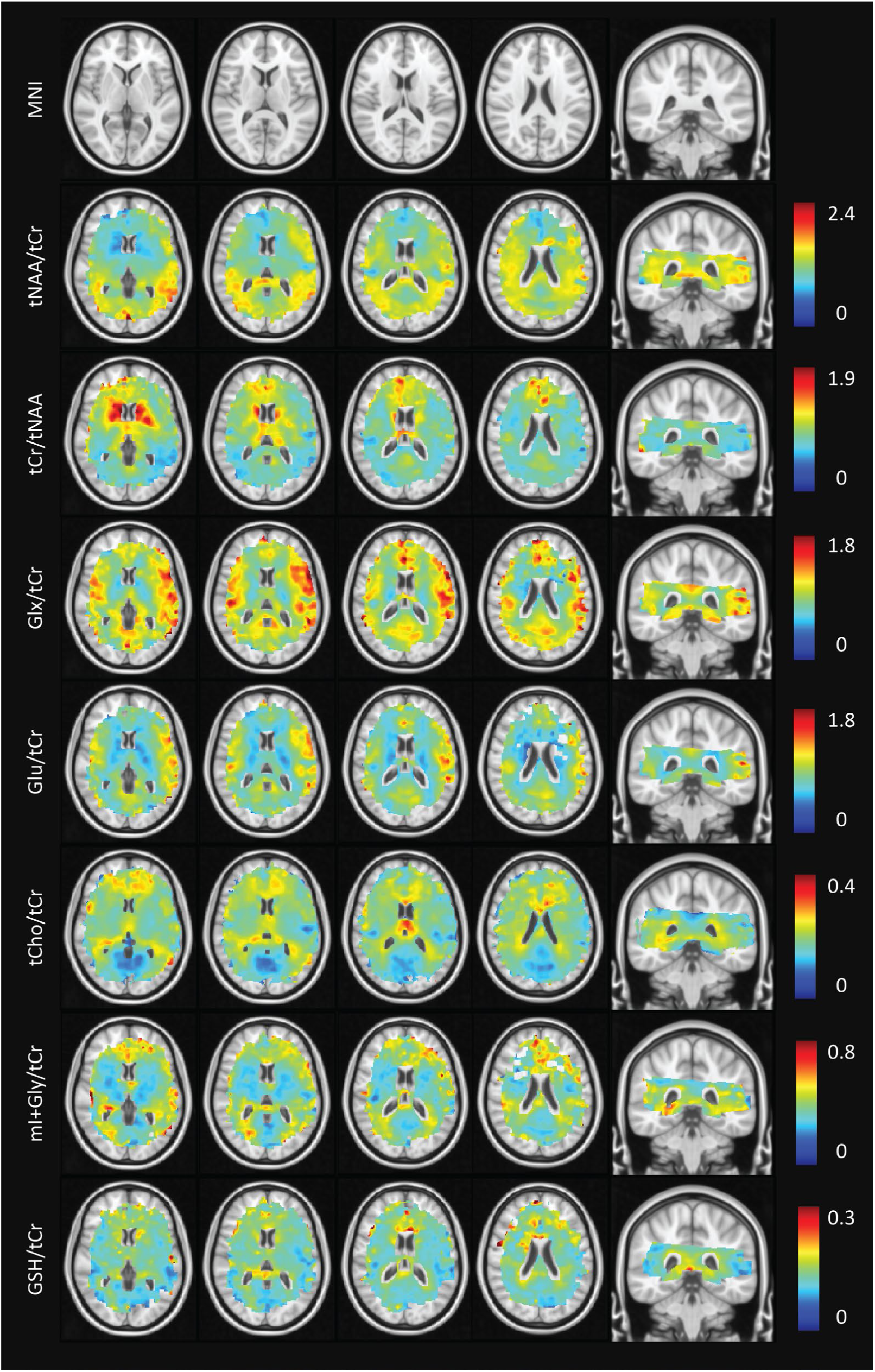
MNI-averaged metabolite maps derived from 23 subjects. Voxels containing less than 3 data points have been removed.

**Figure 6:**
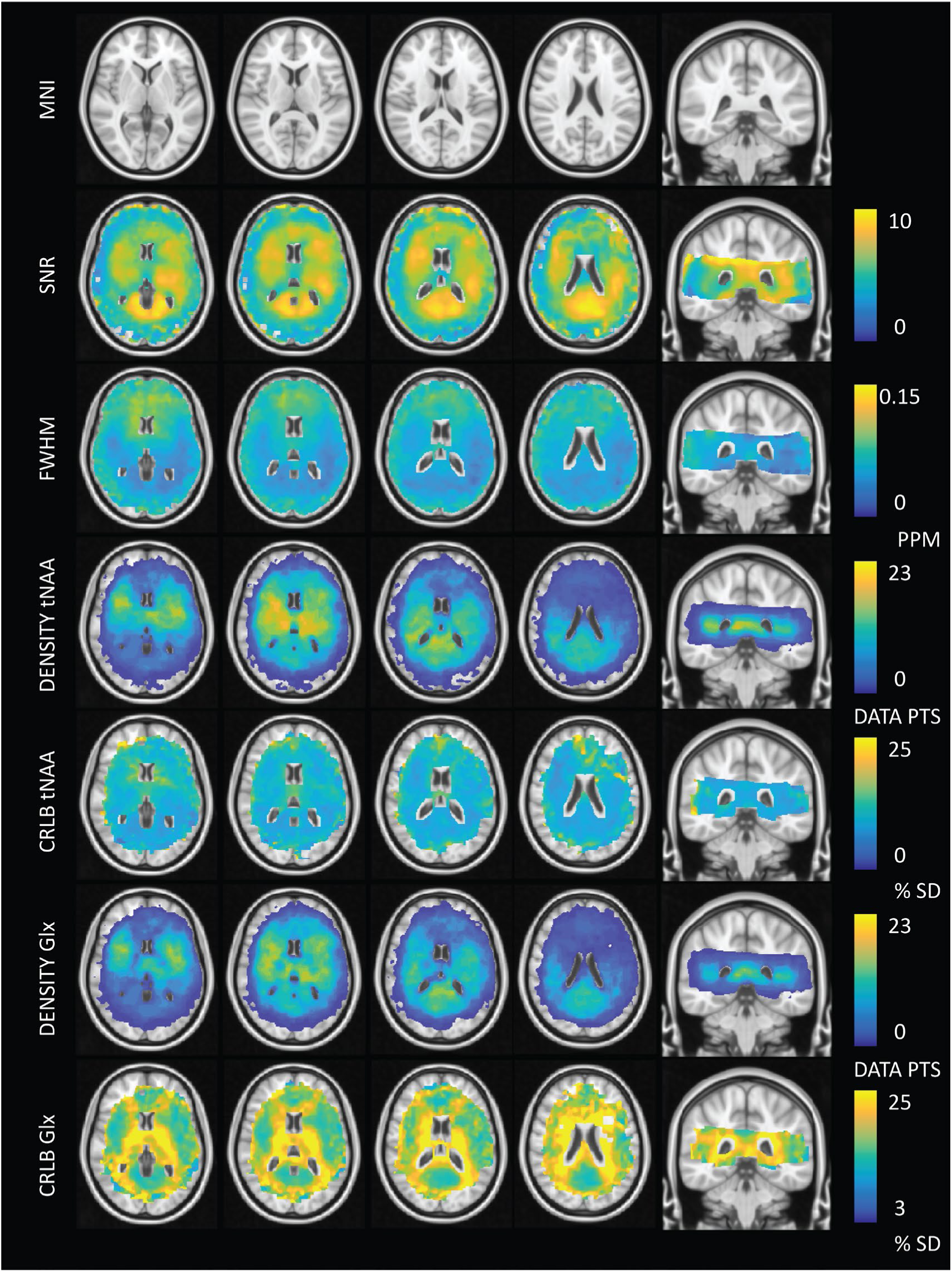
QA maps averaged across 23 subjects. The SNR and FWHM data are output for each individual LCModel fit and show an inverse relationship between signal intensity and linewidth as expected. SNR values less than 3 and FWHM values greater than 0.15 ppm have been filtered out. Density maps for tNAA and Glx are provided. In general, data density is high in central regions of the brain at the location of sub-cortical GM. Corresponding CRLB maps are provided illustrating regions of variable fit confidence. This is notable for Glx, where the CRLB is higher within WM regions. This is in agreement with the fact that Glx is approximately 50% less abundant in white versus gray matter. Lower peak SNR leads to increased fit uncertainty and higher CRLB. CRLB values greater than 50% have been removed.

## 4.0 DISCUSSION

One of the major promises of MRSI is the ability to detect physiological changes in disease in the absence of structural/anatomical markers. While single voxel spectroscopic measurements might be considered the Gold Standard in terms of reliability of data, evolving metabolic mapping techniques provide opportunities for evaluating the metabolic state of the entire brain. Our focus on artefact control, partial volume correction, QA filtration, and data aggregation from multiple subjects has permitted mapping of metabolite ratios metabolites such as tNAA, Glx and tCho at deeper regions of the brain adjacent to the ventricles; areas of interest for studying metabolic alterations linked to psychiatric disorders [27, 47].

An MNI-registered metabolic brain atlas based on pooled MRSI data was first presented by Maudsley et al. [48] who mapped distributions of certain human neuro-chemicals using 3T-EPSI acquisitions. Our work diverges in several important aspects. The use of a crusher coil for direct lipid suppression, instead of post-processing or inversion methods for lipid signal removal, meant that we could acquire MRSI data without the risk of manipulating metabolite signals adjacent to lipid resonances, or residual lipid contamination due to imperfect inversion pulses. Furthermore, in our approach the signals of lactate and macromolecules stay intact and can be taken into consideration during metabolite fitting. Another key difference is our use of the co-registered high-resolution anatomical information for partial volume correction. While partial volume correction of spectroscopic data is not a novel concept, a large majority studies correct for only CSF, whereas we consider tissue specific signal contributions arising from WM and GM at 1mm isotropic resolution. Finally, the higher SNR at 7T has allowed us to generate voxel-wise maps of low signal resonances such as Glu, where groundwork work performed at 3T relied on ROI-based analysis for the derivation of similar information [42].

Our results complement contemporary UHF MRSI reports that tend to focus on brain regions located above the ventricles[2, 3, 49]; likely as a way to reduce sensitivity artefacts stemming from inhomogeneous B_0_ or unsuppressed water while showing the potential of new techniques. Aside from acquisition at optimal brain regions, these innovations can be showcased by optimizing data quality through increased SNR using longer TR, removing (or excluding during metabolite fitting) lipid signals between 0-1.8ppm, or acquiring at higher spatial resolutions at the cost of SNR and scan-time. Our use of tailored water suppression pulses and a crusher coil in combination with a short TE/TR acquisition allowed us to obtain reasonably high resolution, lipid-suppressed MRSI data within 11 minutes. Clinically acceptable scan-times are paramount for widespread adoption of MRSI due to the general requirement of additional anatomical and functional imaging in a typical patient scan-slot. Current efforts to offset scan-time using fast MRSI techniques, show great potential and include accelerated data acquisition via echo planar readout (EPSI) gradients [50, 51] or alternate k-space sampling strategies combined with advanced reconstruction methods [19, 52]. The advantages and limitations of various fast MRSI techniques have been thoroughly reviewed by Shankar et. al [53].

We improve upon previously reported MRSI data registration methods through the use of the high resolution ‘slice’ reconstruction as a surrogate for the MRSI data (figure 2). This reduced mis-registration errors and minimized additional uncertainties relating to inter-subject variability [45]. A consequence of spatially averaging metabolite maps at the resolution of the 1mm MNI152 atlas was an apparent increase in signal contrast between brain regions; particularly in striatal/thalamic regions of the brain where data density was high. This is a point of significance for future group-level comparisons seeking to identify regional differences in small brain structures. A possible drawback of our approach is that we cannot take into account subject motion occurring between the T1w and MRSI scans.

### Considerations

The balance between over-crushing of tissues and under-crushing lipids is a delicate one. Our acquisition protocol was guided by the preference towards reliable signal and this often meant that some tissue signals were over-crushed. This was exacerbated by factors relating to asymmetries in the crushing field (figure 3 and 4) or subject head positioning within the lipid suppression coil itself. Future coil design iterations will focus on greater control of the lipid suppression field through modular design and the use of novel winding patters [54]. Reconstruction [20] and post-processing based [55] lipid removal methods can provide alternative pathways for lipid suppression while conserving MRS signals at the edge of the cortex; however, these methods may have drawbacks relating to signal removal of nearby metabolite resonances [56], or overlapping lactate or macromolecular signals (i.e. around 1.2 and 1.4 ppm). Secondly, while it is advised to include subject specific MM signals into fitting protocols for optimal signal quantification [57], scan-time constraints and the need to generate a new basis per subject left this approach unfeasible. Considering the low resolution of the double inversion-recovery (DIR) acquisitions (30 × 30 × 10mm^3^) along with reports that a general MM baseline is suitable for GM and WM at 7T [58, 59], it was concluded that high quality data from a single volunteer was a reasonable compromise. In line with current literature, we chose to use tCr as an internal reference [60]. Using ratios divides out larger variations associated with inhomogeneous B_1_ transmit/receive fields and coil loading. An important caveat specific to ultra-high field studies is that the inhomogeneity of the excitation pulse across brain regions can lead to variable T1 weightings between metabolites. Absolute quantitation using the water reference along with using additional B_1_ correction steps during post-processing (requiring B_1_ mapping scans) could further improve the accuracy of our reported metabolite ratios. A further source of error may arise under conditions where metabolite peak heights show strong B_1_ dependent T1 weightings. While we corrected for bulk T1 effects per metabolite, we were not able to correct for inhomogeneous relaxation constants between hydrogen-containing molecular subgroups or between different brain regions [61]. This is an issue that is further compounded when considering combined metabolites (tCho, tNAA, Glx or tCr).

On a related note, normalization to tCr assumes a homogeneous distribution across brain regions and can therefore introduce errors under circumstances where tCr levels in the brain have changed due to disease. Disease-related changes in metabolite distributions may also negatively impact the accuracy of metabolite signal scaling during partial volume correction since we currently assume that metabolite ratios for pure tissues remain stable. For this reason, caution is advised if using partial volume correction for areas in which structural changes have occurred (for example, brain tumors or multiple-sclerosis lesions). In general, our measured metabolite values were lower than those reported in literature. Possible sources for these differences are: 1) we include MM signals that overlap with metabolite resonances; 2) we perform T1 and partial volume correction for GM and WM tissue contributions, which is uncommon for MRSI studies. In some cases the correction for the T1 of tCr reduces metabolite ratio values; 3) we use a relatively short TR of 300ms that may introduce T1 weightings particularly for metabolites with long T1s. Based on our flip angle and repetition time, our sequence was optimized to give the highest SNR per unit time for metabolites with T1 values in the range of 1500 to 2000 ms; 4) differences in metabolite fitting software or LCModel control file parameters are known to have a significant impact on peak fitting results [62].

## 5.0 CONCLUSION

In this work we present *in-vivo* metabolite levels for central brain structures obtained by combining several methodological advances, including a crusher coil to suppress the high intensity extracranial lipid signals, advanced reconstruction techniques, partial volume correction based on high-resolution anatomical images and normalization to a standard space. Our acquisition and processing pipeline facilitates group level 7T MRSI analysis and provides an infrastructure to which data from ongoing and future studies using comparable acquisition techniques can be easily added. Given the spatial coverage, values reported herein may be of interest for future studies investigating WM and sub-cortical GM physiology. Registration of MRSI data to standard space opens the possibility to compare and correlate 7T MRSI metabolic data with any parameters normalized into the same space.

## Acknowledgements

This research was supported by a Brain and Behavior Research Foundation NARSAD Young Investigator Award (Christiaan Vinkers, 24074). The authors would like to acknowledge Philippe Cornelisse for his assistance in data acquisition.

## Data Availability Statement

The data that support the findings of this study are available from the corresponding author upon reasonable request.

