## supplemental for "Lipid suppressed and tissue-fraction corrected metabolic distributions in human central brain structures using 2D ^1^H Magnetic Resonance Spectroscopic Imaging at 7 tesla"

**Supplementary figure 1**

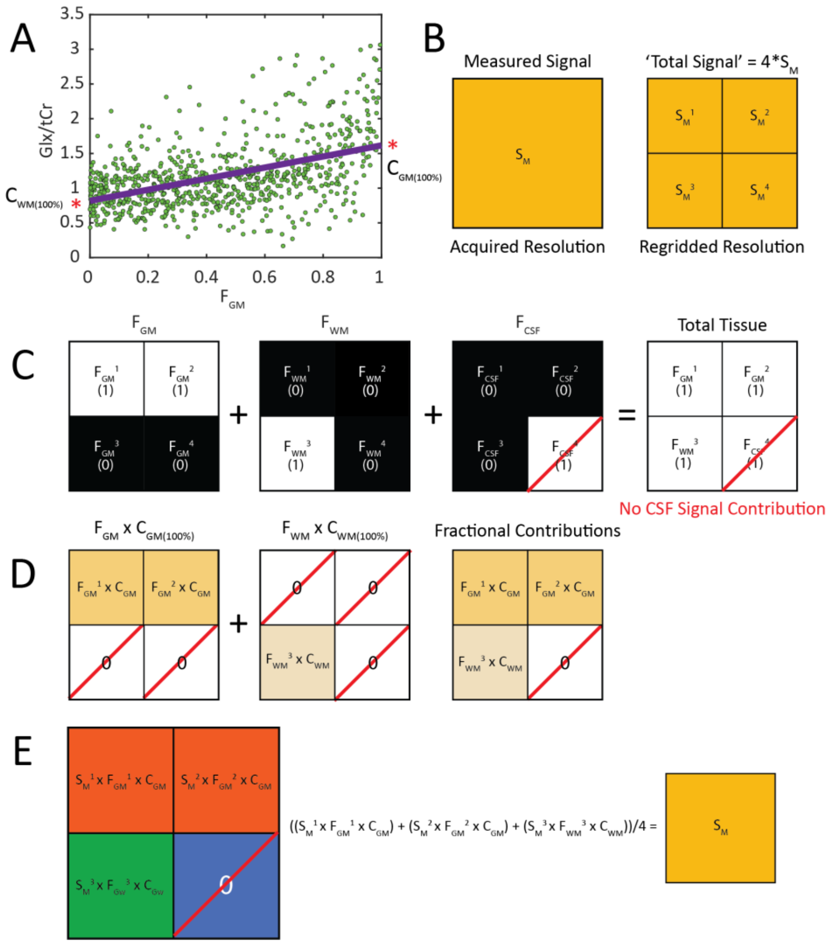

**Graphical representation of partial volume correction**

Figure 1A: scatter plots of normalized GM fraction (GM/GM+WM) as a function of Glx/tCr values across a single subject. Based on these plots, pure tissue contributions for each metabolite can be extrapolated on a per-subject basis. 1B: In this representation, the resolution of the MRSI voxel has been up-sampled by a factor 4 (actual data is up-sampled by a factor 250). Each smaller voxel is attributed with the same value as the source voxel. 1C-D: PSF-corrected partial volume segmentations (manuscript figure 2B) provide the tissue fractions that are multiplied with the pure tissue metabolite values derived from 1A to generate fractional contribution maps accounting for tissue content and pure tissue signal contributions. 1E: Fractional contributions are then multiplied by the initial signal of the source voxel – that has been attributed to each smaller voxel in 1B. The sum of all contributions divided by the total number of voxels will yield a signal value equal to that of the original source voxel. In this way, the average metabolite within the volume occupied by the original source voxel remains unchanged after PVC; however the signal is redistributed in a way that represents the actual tissue distribution within said voxel.

A consideration regarding the PVC described here is that linear regression used to derive the pure tissue concentrations can be weighted by outlier voxels or voxels with altered metabolite signals due to pathology. Errors in the estimation of the pure tissue fractions will propagate when calculating the fractional contributions described in figure 1D/E. Nevertheless, corrected voxel values are based on the measured values meaning that voxels containing low/high values will remain comparatively low/high after correction. Although cumbersome, a possible approach to minimize error propagation might be to consider separate brain regions independently; for example, a unique regression line for deep GM nuclei versus white matter or pathological regions. However, such an approach would come with challenges in dealing with border regions and too few voxels for regression analysis.

### Supplementary figure 2

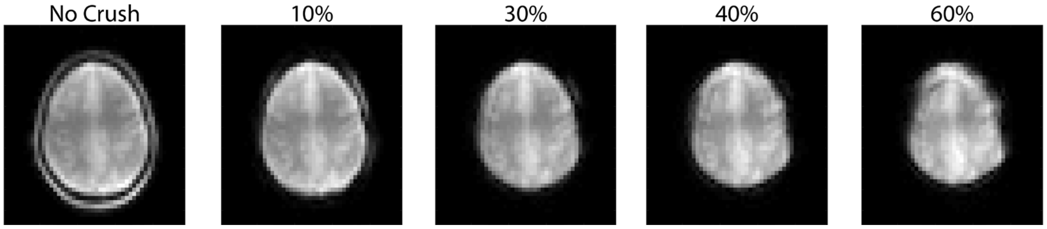

**Low resolution anatomical scans used for crushing field calibration**

Figure 2: The current amplitude to drive the crusher coil was determined empirically for each volunteer. Study Participants were recruited externally and data acquisition time was limited to a 45 minute scan slot. For MRSI data, a low resolution anatomical scan was performed with the amplifier initially set to between 60-80% of its maximum current. The scan was designed to closely mimic the MRSI scan in order to estimate crushing field effectiveness.

Scan parameters: 2D Fast Field Echo sequence, TE/TR = 3/300 ms, FOV = 220 x 200 x 10 mm3, resolution = 4.58 x 4.58 x 10 mm3, scan duration 15 s, flip angle = 1 degree

Generating the scan and updating the crusher coil amplifier settings took approximately 2 minutes. Based on the residual lipid signal, the amplifier strength was adapted to ensure maximal lipid suppression while taking into consideration the amount of tissue signal that was being crushed. This process was repeated for each slice, since crushing fields varied with position and head shape (for example, the top of the head is more round than central regions of the head from which most of our data were acquired). Due to scan time constraints, a maximum of 2 iterations could be performed per slice. For the interested reader, it is important to not that different head shapes along with the location of interest can strongly influence the current amplitude required for optimal lipid suppression. For example, a 60% amplitude may be acceptable for a smaller head size, while the same current could result in significant tissue signal loss when applied to a larger head size. Furthermore, the higher one goes in the brain, the larger the additional effect of coil windings located at the top of the crusher coil’s dome shape.

**Supplementary figure 3**

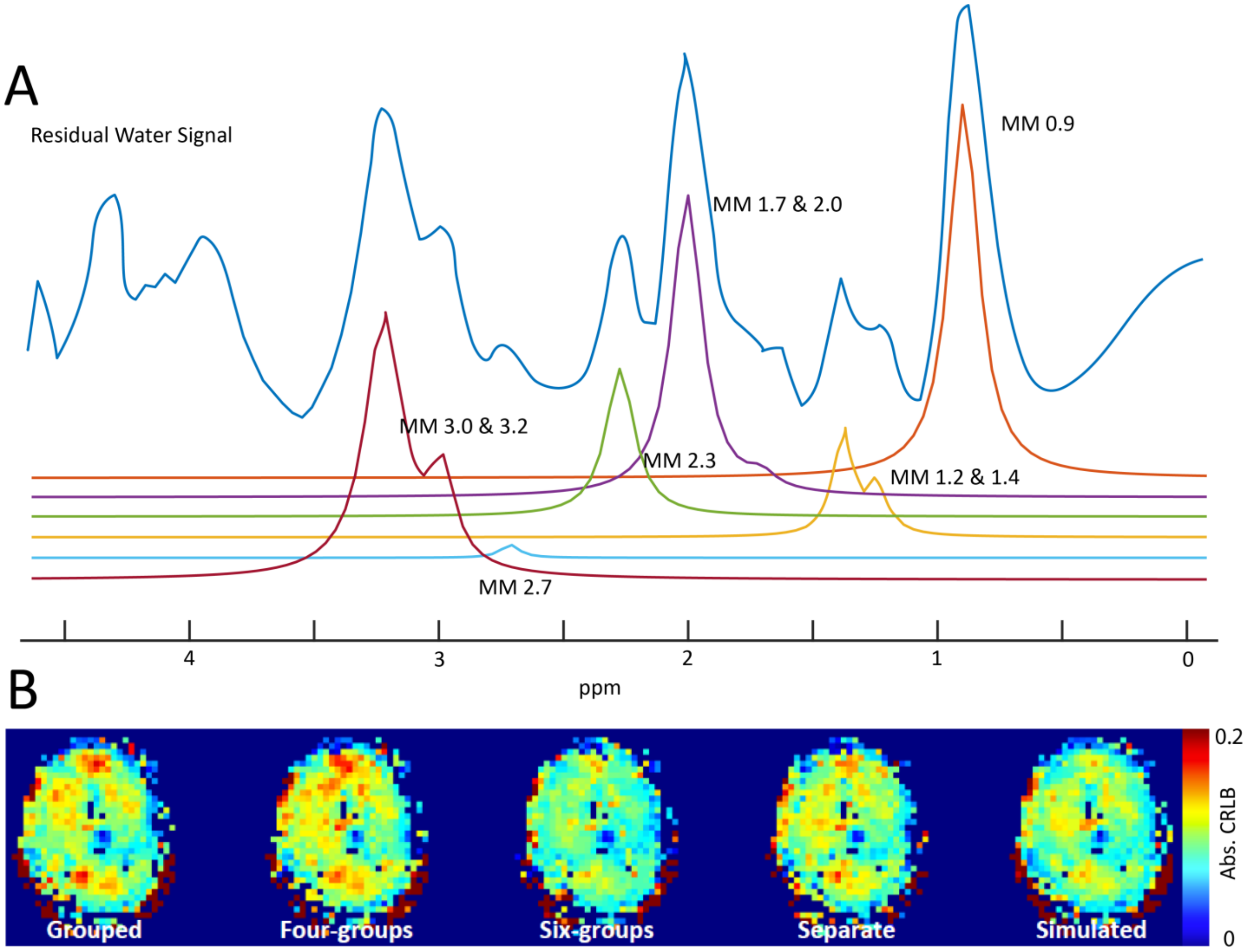

**Macromolecule (MM) lines used in the LCModel basis set**

Figure 3A: the measured MM signal is shown in blue. This signal was decomposed into a series of Gaussian functions representing individual MM peaks as shown. Initially, several different variations of MM basis functions were added to the LCModel basis set including: fully grouped, four groups (3.0+3.2+2.7, 2.3+1.7+2.0, 1.2+1.4, 0.9), six groups (3.0+3.2,2.7, 2.3,1.7+2.0, 1.2+1.4, 0.9), separate and simulated. Selection of the preferred MM grouping was made based on a minimization of CRLB values of the glutamate fit since low CRLB are indicative of a higher degree of fit certainty. Figure 3B shows an example of glutamate CRLBs for each condition in a single subject. Grouped MM are mutually constrained in that they are treated as a single basis line in the LCModel fit. No hard constraints were imposed on the MM basis functions.

**Supplementary figure 4**

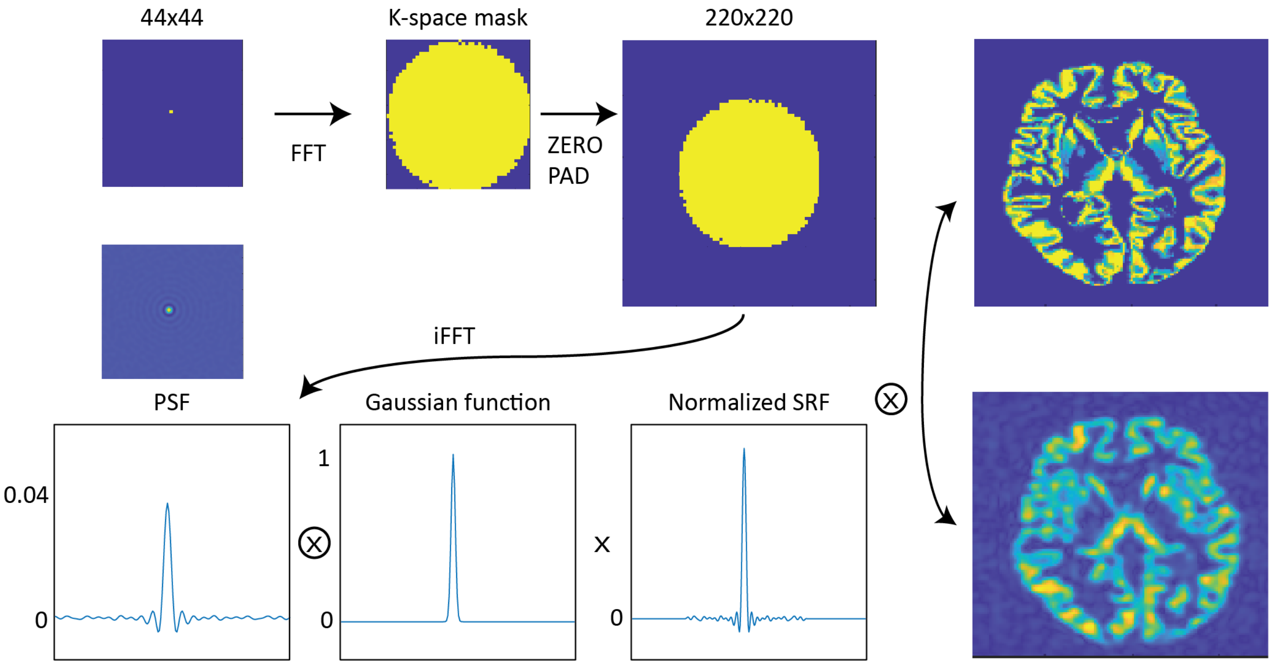

**Conceptual outline for generation of the point-spread function used for anatomical data during partial volume correction**

Figure 4: In order to properly correct for tissue contributions, it is important to adapt the anatomical segmentations such that they better represent what is 'seen' from the perspective of the MRSI voxel. Due to the lower resolution acquisition grid of the MRSI scan, voxel bleeding from tissues located outside of the nominal voxel can occur. These contributions must be taken into account.

The first step in generating the PSF was to create a single voxel in a 44x44 grid. The resulting image was Fourier transformed and a k-space mask based on the k-space shutter used during acquisition was applied. The masked k-space was then zero padded to have an acquisition matrix the same size as that of the anatomical segmentations. Next an inverse Fourier transform was applied to generate the PSF. During reconstruction of the MRSI data, a Gaussian response function was applied as described in: *Kirchner, T., et al., Reduction of voxel bleeding in highly accelerated parallel (1) H MRSI by direct control of the spatial response function. Magn Reson Med, 2015. 73(2): p. 469-80.* The PSF was also convolved with the same response function and normalized. The final PSF (or spatial response function (SRF)) was convolved with each anatomical segmentation to introduce the blurring as shown in the right panel. Metabolic maps were resampled to the resolution of the PSF-convolved anatomical segmentations in order to facilitate the partial volume correction method outlined in supplementary figure 1.

**Supplementary table 1**

| Metabolite | T1 WM (s) | T1 GM (s) |
|---|---|---|
| Cr | 1.78 | 1.74 |
| Cho | 1.32 | 1.51 |
| mIGly | 1.19 | 1.28 |
| tCr | 1.78 | 1.74 |
| tNAA | 1.9 | 1.83 |
| tCho | 1.32 | 1.51 |
| Glx | 1.75 | 1.61 |
| Gln | 1.74 | 1.64 |
| Glu | 1.75 | 1.61 |
| NAA | 1.9 | 1.83 |
| NAAG | 0.94 | 1.21 |
| GPC | 1.32 | 1.51 |
| PC | 1.32 | 1.51 |
| PCr | 1.78 | 1.74 |
| Tau | 2.09 | 2.15 |
| mI+Gly | 1.19 | 1.28 |
| GSH | 1.06 | 1.14 |
| Macromolecules | 0.42 | 1.14 |

Tissue specific values used for T1 correction of metabolic maps. Taken from Xin et. al, MRM 2013

**Supplementary table 2**

Table 1

| Average percentage of voxels removed upon filtration by FWHM & SNR maps | |
|---|---|
| FWHM | 15 ± 3 |
| SNR | 22 ± 8 |

Average percentage of metabolite voxels removed

| Metabolite | All Filters | Component from CRLB + Lipid |
|---|---|---|
| tCr | 56 ± 10 | 7 |
| tCho | 51 ± 10 | 12 |
| GSH | 45 ± 12 | 18 |
| Glx | 51 ± 11 | 12 |
| Glu | 48 ± 12 | 15 |
| tNAA | 53 ± 10 | 10 |
| mI + Gly | 45 ± 11 | 18 |

The percentage of removed voxels was calculated on an individual per metabolite map basis and is influenced by factors such as brain masking and CSF fraction. For instance, slices lower in the brain will lose significant numbers of voxels when they contain ventricles. Metabolite maps were first filtered based on SNR and FWHM values output from LCModel. Next, a CRLB threshold of 50% was applied to the tCR map and Lipid map. The result was binarized to create a lipid filter. This filter, along with the individual CRLB maps (binarized and including only voxels with CRLB < 50%) were applied to each metabolite map respectively. Average maps generated from individual filtered metabolite maps display larger metabolite coverage than what might be expected based on filtration metrics since not all voxels are filtered in the same way for all subjects. This demonstrates one of the benefits associated with using a common reference space for MRSI data.

**Supplementary data: LCModel control parameters**

sddegp= 7
ppmst= 4.0
ppmend= 0.2
nunfil= 519
nomit= 8
neach= 50
ndslic= 1
ndrows= 44
ndcols= 44
ltable= 7
lps= 8
lcsv= 11
lcoraw= 10
lcoord= 9
islice= 1
irowst= 5
irowen= 40
icolst= 5
icolen= 40
hzpppm= 3.0000e+02
echot= 2.50
dkntmn= 0.3
dgppmx= 20
dgppmn= -20
deltat= 3.333e-04
degppm= 0
chomit(8)= 'MM17'
chomit(7)= 'MM20'
chomit(6)= 'MM14'
chomit(5)= 'MM12'
chomit(4)= 'MM09'
chomit(3)= 'PE'
chomit(2)= 'Ace'
chomit(1)= 'Ala'
$END
